# Computational simulation of the reactive oxygen species and redox network in the regulation of chloroplast metabolism

**DOI:** 10.1101/638437

**Authors:** Melanie Gerken, Sergej Kakorin, Kamel Chibani, Karl-Josef Dietz

## Abstract

Cells contain a thiol redox regulatory network to coordinate metabolic and developmental activities with exogenous and endogenous cues. This network controls the redox state and activity of many target proteins. Electrons are fed into the network from metabolism and reach the target proteins via redox transmitters such as thioredoxin (TRX) and NADPH-dependent thioredoxin reductases (NTR). Electrons are drained from the network by reactive oxygen species (ROS) through thiol peroxidases, e.g., peroxiredoxins (PRX). Mathematical modeling promises access to quantitative understanding of the network function and was implemented for the photosynthesizing chloroplast by using published kinetic parameters combined with fitting to known biochemical data. Two networks were assembled, namely the ferredoxin (FDX), FDX-dependent TRX reductase (FTR), TRX, fructose-1,6-bisphosphatase pathway with 2-cysteine PRX/ROS as oxidant, and separately the FDX, FDX-dependent NADP reductase (FNR), NADPH, NTRC-pathway for 2-CysPRX reduction. Combining both modules allowed drawing several important conclusions of network performance. The resting H_2_O_2_ concentration was estimated to be about 30 nM in the chloroplast stroma. The electron flow to metabolism exceeds that into thiol regulation of FBPase more than 7000-fold under physiological conditions. The electron flow from NTRC to 2-CysPRX is about 5.46-times more efficient than that from TRX-f1 to 2-CysPRX. Under severe stress (30 μM H_2_O_2_) the ratio of electron flow to the thiol network relative to metabolism sinks to 1:251 whereas the ratio of electron flow from NTRC to 2-CysPRX and TRX-f1 to 2-CysPRX rises up to 1:80. Thus, the simulation provides clues on experimentally inaccessible parameters and describes the functional state of the chloroplast thiol regulatory network.

**Authors summary:** The state of the thiol redox regulatory network is a fundamental feature of all cells and determines metabolic and developmental processes. However, only some parameters are quantifiable in experiments. This paper establishes partial mathematical models which enable simulation of electron flows through the regulatory system. This in turn allows for estimating rates and states of components of the network and to tentatively address previously unknown parameters such as the resting hydrogen peroxide levels or the expenditure of reductive power for regulation relative to metabolism. The establishment of such models for simulating the performance and dynamics of the redox regulatory network is of significance not only for photosynthesis but also, e.g., in bacterial and animal cells exposed to environmental stress or pathological disorders.

## INTRODUCTION

Reduction-oxidation reactions drive life. In aerobic metabolism, electrons from reduced compounds pass on to oxygen to produce water and ATP. Photosynthesis exploits light energy and reverses this oxidation process by water splitting, liberation of O_2_ and reduction of CO_2_, NO_3_” and SO_4_^2−^ to carbohydrates, amines and sulfhydryl compounds. A decisive role is played by ferredoxin (FDX) which functions as hub of electron distribution accepting electrons from photosystem I and donating them in particular to FDX-dependent NADP reductase (FNR), FDX-dependent nitrite reductase (NIR), FDX-dependent sulfite reductase (SIR), FDX-dependent glutamate oxoglutarate aminotransferase (GOGAT), FDX-dependent thioredoxin reductase (FTR) and to O_2_ in the Mehler reaction [1]. Considering the elemental composition of a typical plant body, C:N:S need to reduced and incorporated at a ratio of roughly 40:8:1. The establishing of this ratio and avoidance of wasteful processes requires fine-tuned regulation of electron flows and metabolism.

The adjustment of metabolic fluxes in the chloroplast to a major extent is controlled by electron flow into the thiol redox regulatory network. Polypeptides switch from an oxidized form with intra-or intermolecular disulfide bridges to a reduced thiol state. TRX and the chloroplast NADPH-dependent TRX reductase C (NTRC) act as electron transmitters in the reduction process. NTRC combines a NADPH-dependent TRX reductase domain with a TRX domain [2]. The TRX complement of Arabidopsis plastids comprises 20 TRX and TRX-like proteins with representatives of the f-, m-, x-, y-, z-group of TRX, TRX-like proteins which include chloroplast drought-induced stress protein of 32 kDa (CDSP32), Lilium1-4 (ACHT1-4) and TRX-like [3]. TRX-f1 and TRX-f2 function in activation of Calvin-Benson-Cycle (CBC) enzymes and γ-subunit of F-ATP synthase [4]. TRX-m1, -m2, -m3 and -m4 are suggested to regulate targets which control the NADPH/NADP ratio [5] which is linked to their ability to efficiently activate the NADPH-dependent malate dehydrogenase [6]. TRX-x, NTRC and TRX-lilium1(ACHT4a) were identified as reductants of the 2-cysteine peroxiredoxin (2-CysPRX) [7–9], and TRX-y1 and –y2 as reductant of PRX-Q [10]. These exemplary studies describe specificity and redundancy for the interaction between TRX-forms and target proteins, as, e.g., comparatively investigated by Collin et al. [7].

Upon transition from dark to light or upon an increase in photosynthetic active radiation (PAR) reductive activation of CBB enzymes via redox sensitive thiols stimulates consumption of NADPH and ATP and coordinates energy provision in the photosynthetic electron transport (PET) chain and energy consumption in metabolic pathways. However it is less understood how once activated enzymes are down-regulated by oxidation. Oxygen and reactive oxygen species (ROS) function as final electron acceptors. ROS generated in the PET react with thiol peroxidases (TPX) with high affinity [11]. Redox transmitters regenerate oxidized TPX. In case of 2-CysPRX, NTRC most efficiently reduces the oxidized form. Other redox transmitters such as TRX-f1, Trx-m1 or Trx-like proteins like CDSP32 also reduce 2-CysPRX at lower rates [7]. The main pathway of TRX reduction targets proteins via FDX and FTR and prevails in strong light. In addition NTRC provides electrons to 2-CysPRX which compensates for the oxidation of 2-CysPRX by PET-produced H_2_O_2_ [2]. The drainage of electrons from other TRXs to oxidized 2-CysPRX may be insignificant under these conditions. This situations changes in darkness were the rate through the PET-driven FDX/TRX-pathway mostly ceases or at lowered photosynthetic active radiation where intermediate flux conditions are established. This balance between oxidation and reduction is suggested to determine the rate of, e.g., the Calvin Benson cycle [12]. While the experimental evidence supports the functionality of this regulatory model, quantitative understanding of the interacting electron fluxes within the network cannot be obtained exclusively from experiments but requires mathematical simulation of the involved major pathways. For this reason this study aimed to first simulate individual electron pathways and then to combine them for predicting crucial parameters of the network inaccessible to experimental determination. Using this approach, it was possible to estimate relative electron fluxes directed into carbon reduction and thiol dependent regulation, and to estimate the rate of H_2_O_2_ production in the chloroplast.

## RESULTS

Ferredoxin (FDX) functions as hub for electron distribution at the donor site of photosystem I. The first mathematical model aimed to simulate electron distribution from TRX-f1 to FBPase and 2-CysPRX in dependence on the H_2_O_2_ concentration (Fig. 1A) and was built on the model presented by Vaseghi et al. [12]. The question asked concerned the efficiency of oxidized 2-CysPRX to compete with reduction of TRX-f1 by FDX-dependent TRX reductase (FTR). The H_2_O_2_ concentration was adjusted to values between 0.3 nM and 10 μM and the steady state redox states of FTR, TRX-f1, FBPase and 2-CysPRX were modelled by kinetic simulation (Fig. 1B-E). The FTR was highly reduced under all conditions and there was only a slight increase from 0.12% to 0.19% oxidation if the H_2_O_2_ rose from 1 nM to 100 nM. Further elevation of H_2_O_2_ had no further effect since the 2-CysPRX turned maximally oxidized at 100 nM and higher H_2_O_2_ concentrations. In the same range, the oxidized form of TRX-f1 reached 50%, while the FBPase was oxidized by 65%.

**FIGURE 1:**
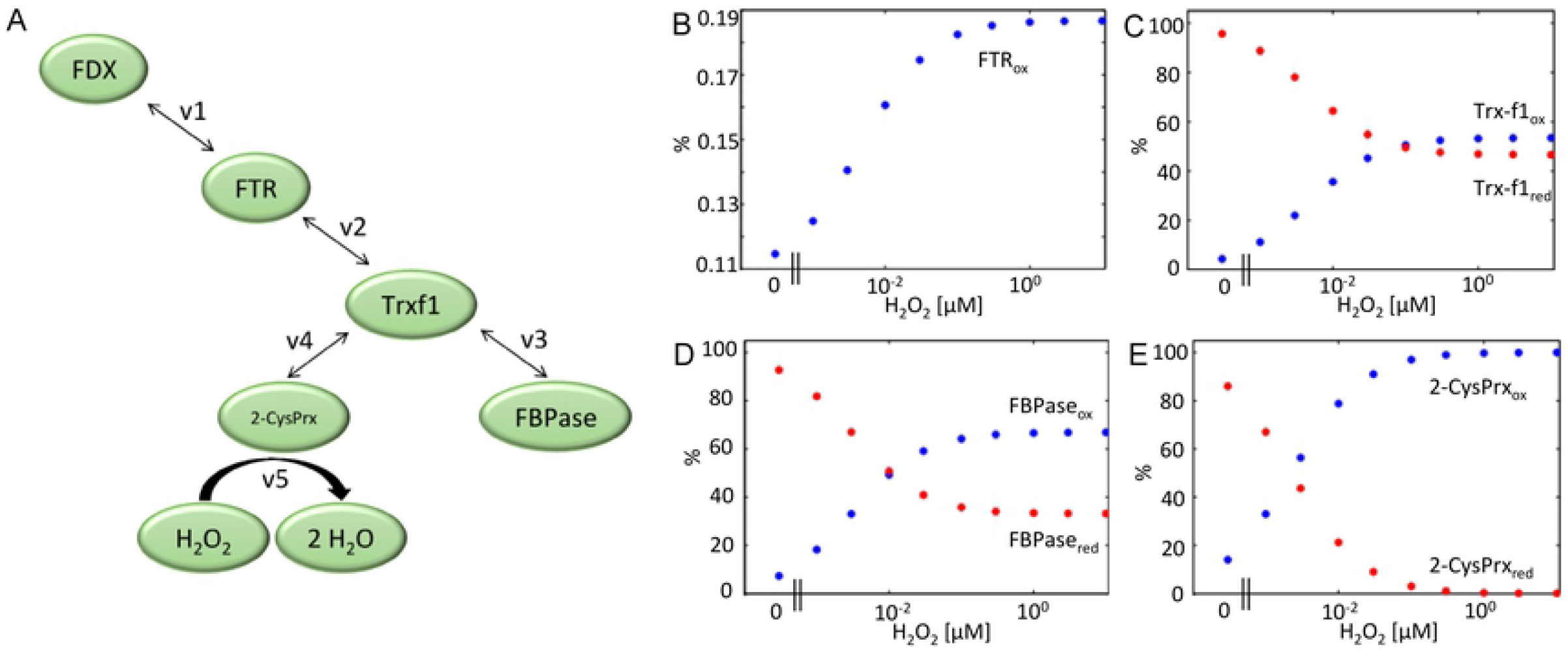
Simulated redox state of FTR-network components in dependence on H_2_O_2_ concentration. (A) Schematic representation of the FTR-network. Electrons are drained from FDX through FTR and TRX-f1 to either FBPase or 2-CysPRX which in turn is oxidized by H_2_O_2_. Each component switches between the reduced and oxidized state. The concentrations were calculated for 1 mg Chl (Supplementary Table 1). FDX was clamped to 50 % reduced state. Starting values of FTR and TRX-f1 were set to 80 % reduced and 20 % oxidized. 2-CysPRX start values for reduced and oxidized form were 35% and 65% [12]. (B-E) Redox states of the network components FTR, TRX-f1, 2-CysPRX and FBPase at varying H_2_O_2_ concentrations as obtained after 3h of simulation in the presence of H_2_O_2_ ranging between 0 and 10 μM.

Fig. 2 depicts the time-dependent changes in redox potential of the sub-network components FTR, TRX-f1, FBPase and 2-CysPRX. In the absence of H_2_O_2_ or at 1 nM, the starting condition shifted to a slightly more reduced state. On the contrary, the FBPase redox potential was poorly affected by increasing the H_2_O_2_ concentration from 1 to 10 μM and even 100 nM already strongly oxidized the TRX-f1 and FBPase proteins. Thus this simple network simulation together with the reported ex vivo redox states of the components allowed us to predict that stromal H_2_O_2_ levels likely range somewhere between 1 and 100 nM.

**FIGURE 2:**
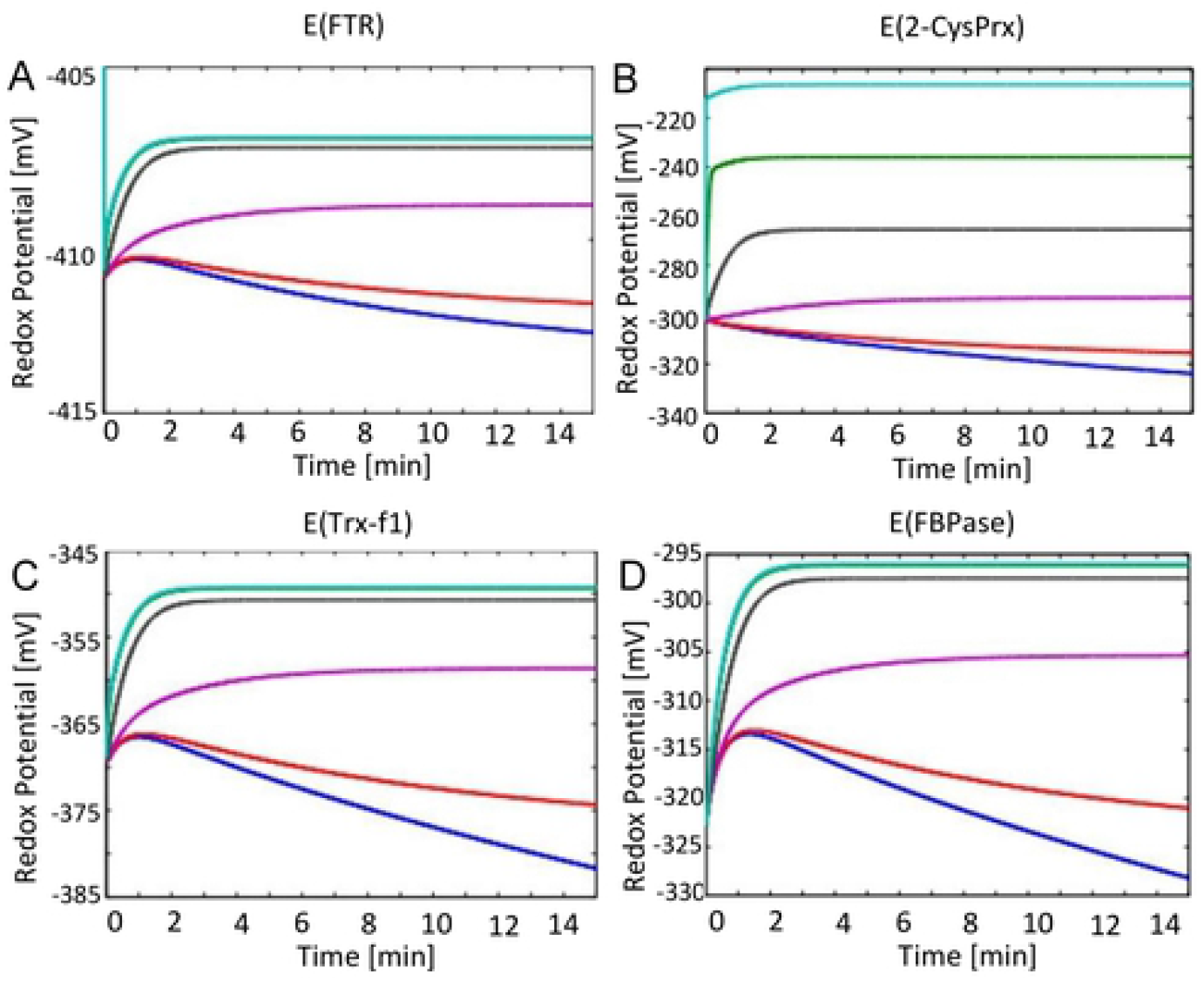
Time-dependent simulation of redox potential changes of FTR-network components. The redox potentials of FDX, FTR, TRX-f1, FBPase, 2-CysPrx were simulated at varying H_2_O_2_ concentrations. Redox potentials were calculated at each time step using the Nernst equation for (A) FTR, (B) 2-CysPRX, (C) TRX-f1 and (D) FBPase. The simulation was run for 15 min for each H_2_O_2_ concentration adjusted to 0 nM (blue), 1 nM (red), 10 nM (magenta), 100 nM (black), 1 μM (green) and 10 μM (cyan).

The second model was constructed to simulate the FNR branch of the network (Fig. 3). Generated NADPH provided electrons to metabolism (v7) or to NTRC for reducing 2-CysPRX. H_2_O_2_ was adjusted to concentrations between 0 and 100 μM. Fig. 3B-E depicts the relative redox forms computed for simulated 3 h which essentially represents the final steady state. The most sensitive component of the network was 2-CysPRX. At about 10 nM H_2_O_2_, the 2-CysPRX was half reduced and half oxidized. NTRC and NADPH responded significantly, here considered as increase in oxidation by at least 10%, when H_2_O_2_ reached a concentration of 1 μM.

**FIGURE 3:**
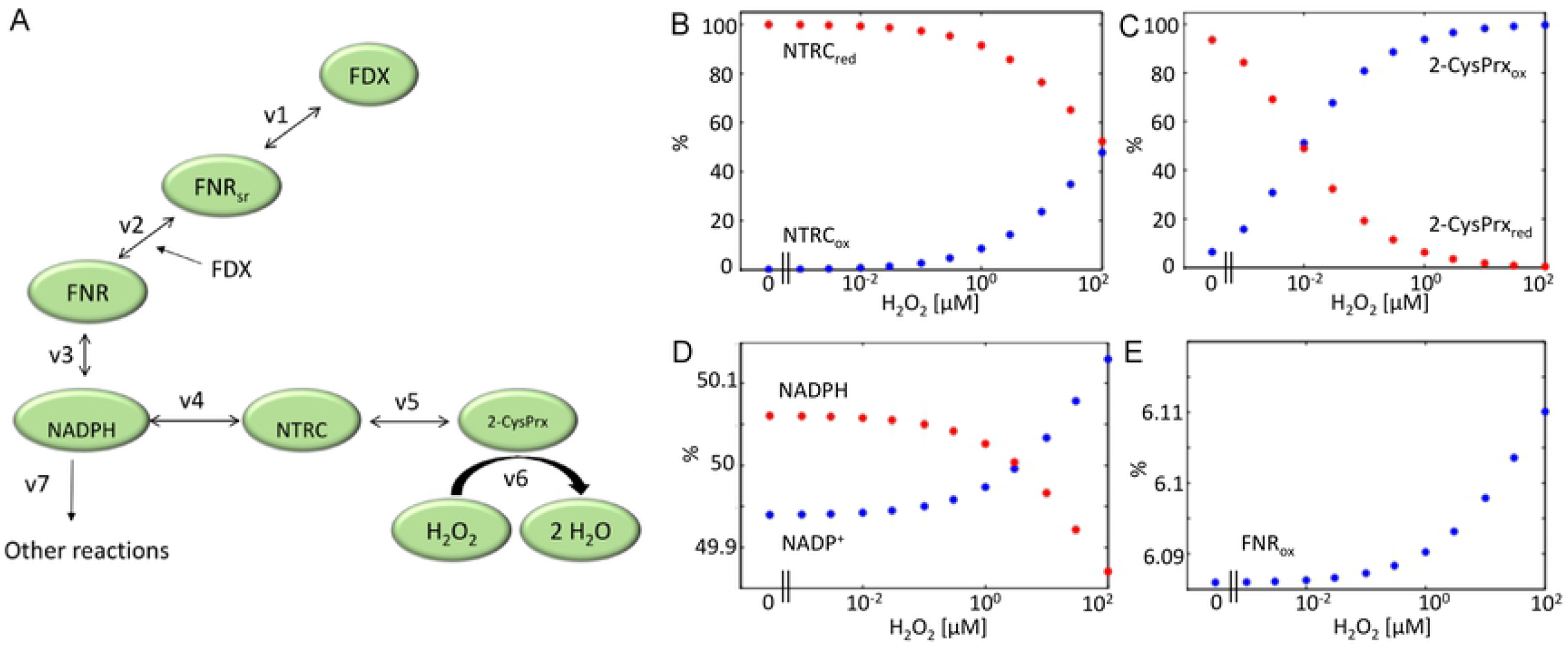
Simulated steady state concentration of FNR-network components at various H_2_O_2_ concentrations. (A) Schematic representation of FNR-network simulated in the second model. Here, electrons passed from FDX through FNR, NADP^+^, NTRC to 2-CysPRX and finally H_2_O_2_. Each component was able to adopt a reduced or oxidized state. FNR is represented in three states in the model; reduced (red), semi reduced (semired) and oxidized (ox). The physiological concentrations were calculated for 1 mg Chl (Suppl. Table 2). FDX was clamped to 50 % reduction. Initial values of FNR_red_ and FNR_semired_ were set to 40 %. All other oxidized forms were initially set to 20 % apart from 2-CysPRX at the starting point with 65% in the oxidized form [12]. The NADPH/NADP^+^ couple was full reduced at t=0. To mimic metabolic NADPH oxidation an additional reaction constant (v7) was added. (B-E) The redox state of the network components (B) NTRC, (C) 2-CysPRX, (D) NADPH, NADP^+^ and (E) FNR_ox_, was simulated for 3h at constant H_2_O_2_ concentrations varying from 0 μM to 100 μM.

The simulation of the FNR-network presented in Fig. 4 focused on the time-dependent changes in redox potentials. The increase of the clamped H_2_O_2_ concentration from 10 (magenta) to 100 nM (black) switched the trend from increased reduction, equivalent to more negative redox potentials, to more oxidation which is equivalent to less negative redox potentials.

**FIGURE 4:**
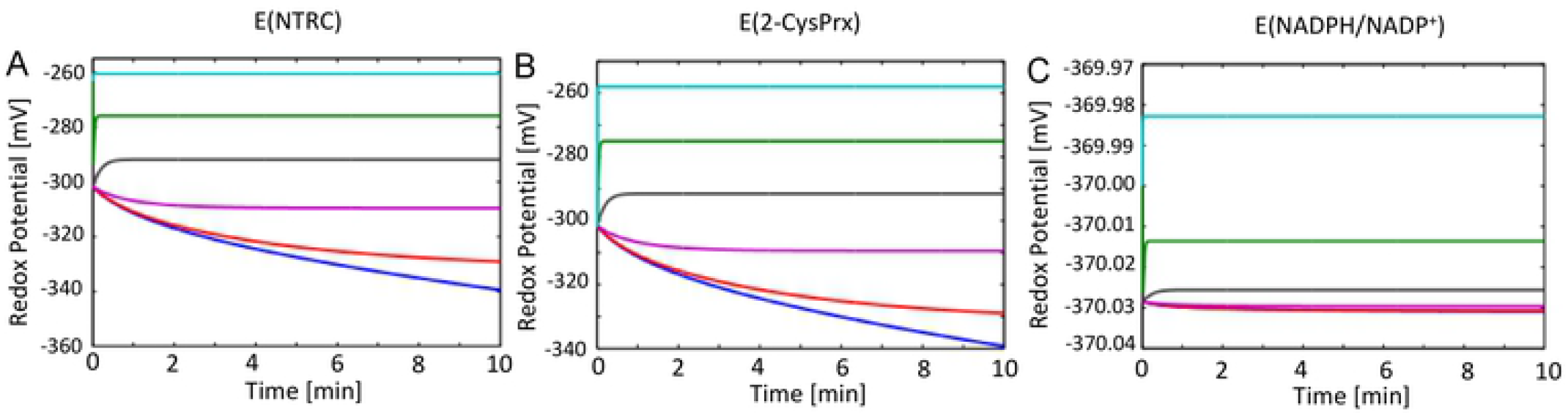
Time-dependent simulation of the redox potentials of the FNR-network components. The redox potentials were simulated for the FNR-network components Fd, FNR, NADPH, NTRC and 2-CysPrx in dependence of the clamped H_2_O_2_ concentration. Redox potentials were calculated at each time step using Nernst equation for (A) NTRC, (B) 2-CysPRX and (C) NADPH/NADP^+^ couple. The simulation was run for 10 minutes at constant H_2_O_2_ concentration of 0 μM (blue), 1 nM (red), 10 nM (magenta), 100 nM (black), 1 μM (green) and 10 μM (cyan).

In the next step, the FTR and FNR networks were combined (Fig. 5A). The H_2_O_2_ concentration was clamped to values between 0 and 100 μM as before and the redox states of the components derived in the approximated steady state after 3h of simulation (Abb. 5B-I). The H_2_O_2_ concentration dependencies of the redox states at first glance were rather similar between the individual and the combined models; however there were some striking differences with likely physiological significance. The TRX-f1 was still more reduced at 100 nM H_2_O_2_ in the combined than in the FTR model. Accordingly, the FBPase remained more reduced in the combined model still being 50% reduced at 100 nM H_2_O_2_ while it was close to half oxidized at 10 nM in the FTR model (cf. Fig. 1 and 5).

**FIGURE 5:**
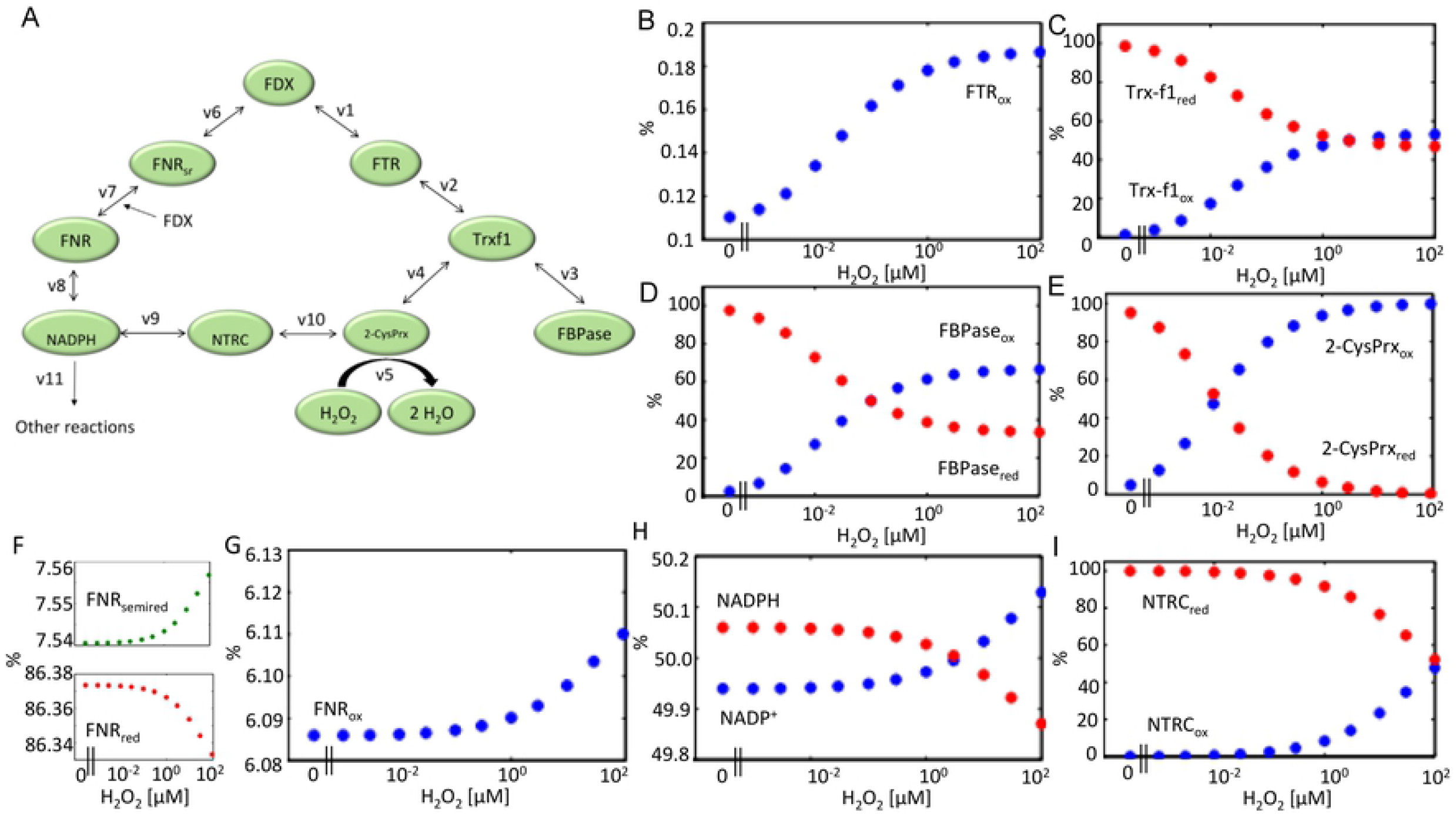
Simulation in the combined model of the redox states of the chloroplast FTR/FNR-network components in the presence of varying H_2_O_2_ concentrations. (A) Schematic representation of the combined FTR/FNR-network model. Electrons from FDX could flow either through the FNR branch to NADP^+^ and NTRC or were transported through the FTR branch to TRX-f1 and FBPase. Thus electrons were transferred to 2-CysPRX and H_2_O_2_ by NTRC and TRX-f1. Each component adopted either a reduced or oxidized state. FNR is represented in three states in the model, the reduced (red), semi reduced (semired) and oxidized (ox) form. The physiological concentrations are calculated for 1 mg chlorophyll (Suppl. Table 3). FDX was clamped to 50 % reduced state. Estimated start values of FNR_red_ and FNR_semired_ were each set to 40 %. The oxidized form was initially set to 20 %. Initial values of NTRC, FTR and TRX-f1 were 80 % reduced and 20 % oxidized. The initial 2-CysPRX values were set to 35% reduced and 65% oxidized form [12]. The NADPH/NADP^+^ couple started from a fully reduced state at t=0. To mimic metabolic NADPH oxidation, the reaction constant v11 was added. (B-E) The redox states of the network components (B) FTR_ox_, (C) TRX-f1, (D) FBPase, (E) 2-CysPRX, (F,G) FNR, (H) NADPH, NADP^+^ and (I) NTRC were simulated for 3h at constant H_2_O_2_ concentrations ranging from 0 μM to 100 μM.

The most striking difference was seen for 2-CysPRX which was half oxidized at 2 nM H_2_O_2_ in the FTR model, but at slightly above 10 nM in the combined model. These important alterations in redox state after introducing the FNR branch witness the importance of the NTRC pathway in reducing 2-CysPRX in line with experimental results such as those published [9, 13]. The time-dependent changes in redox states of the network components (Fig. 6) confirmed the critical range of the H_2_O_2_ concentration needed for stable redox states as also measurable *ex vivo*. Thus at 10 nM H_2_O_2_ in the combined model, there was a trend towards higher reduction, while clamping the H_2_O_2_ concentration to 100 nM reversed the trend toward higher oxidation of the network components.

**FIGURE 6:**
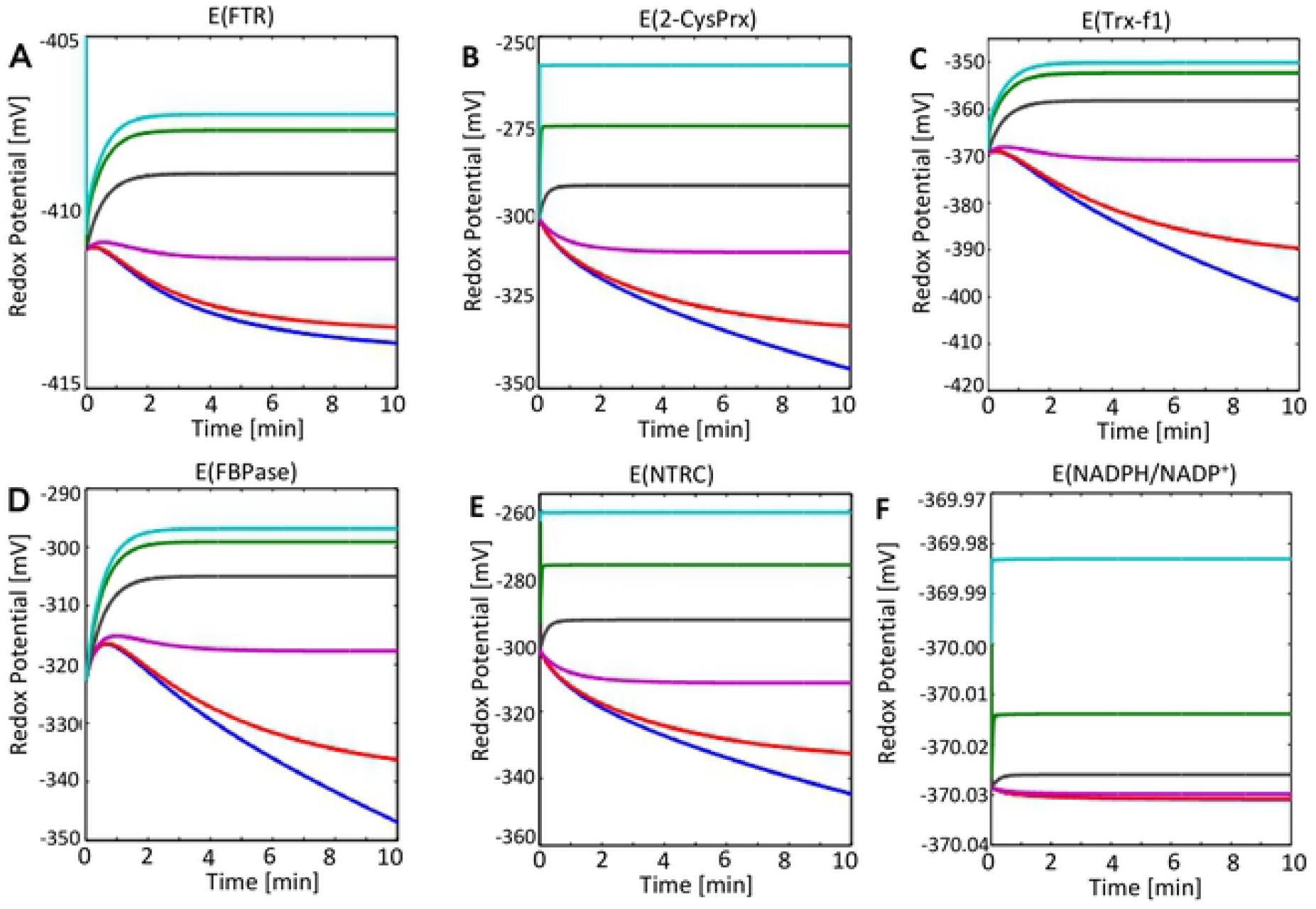
Time-dependent simulation of redox potentials of FTR/FNR-network components. The redox potentials of FTR/FNR-network components were simulated and included FDX, FNR, NADPH, NTRC, FTR, TRX-f1, FBPase and 2-CysPrx. Redox potentials were calculated at each time step using the Nernst equation for (A) FTR, (B) 2-CysPrx, (C) Trx-f, (D) FBPase, (E) NTRC and (F) NADPH/NADP^+^ couple. The simulation was run for 10 min at constant H_2_O_2_ concentrations of 0 μM (blue), 1 nM (red), 10 nM (magenta), 100 nM (black), 1 μM (green) and 10 μM (cyan).

The combined FTR/FNR-model allowed for estimating relative rates of electron drainage at competing branching points of the network and provided answers to the critical questions raised above. The first question addressed the estimation of the resting H_2_O_2_ concentration in the stroma *in vivo*. Several experimental studies have shown that the oxidized fraction of 2-CysPRX exceeds that of the reduced fraction, e.g., [12] determined the ratio of oxidized to reduced forms to 65%:35%. Thus we asked our model at which clamped H_2_O_2_ concentration this particular ratio is realized (Suppl. Table 3). The ratio of 65%:35% was established at 30 nM H_2_O_2_.

The second question concerned the ratio of electron flows from NADPH into metabolism (v11) and NTRC reduction (v9) assuming that only 2-CysPRX acts as electron sink (Fig. 7). For answering this question it was assumed that the H_2_O_2_ concentration in the resting state is close to 30 nM and then the simulated rate constants v9 and v11 and their ratios were computed (Suppl. Table 4). In this scenario, the electron flow into metabolism exceeded that into NTRC-dependent regeneration of 2-CysPRX by a factor of 7234. The rate of regulatory electron flow reached only 0. 14‰ of metabolic reduction. This value increased with increasing H_2_O_2_ in the simulation but did not exceed 5‰ even in the presence of 100 μM H_2_O_2_.

**FIGURE 7:**
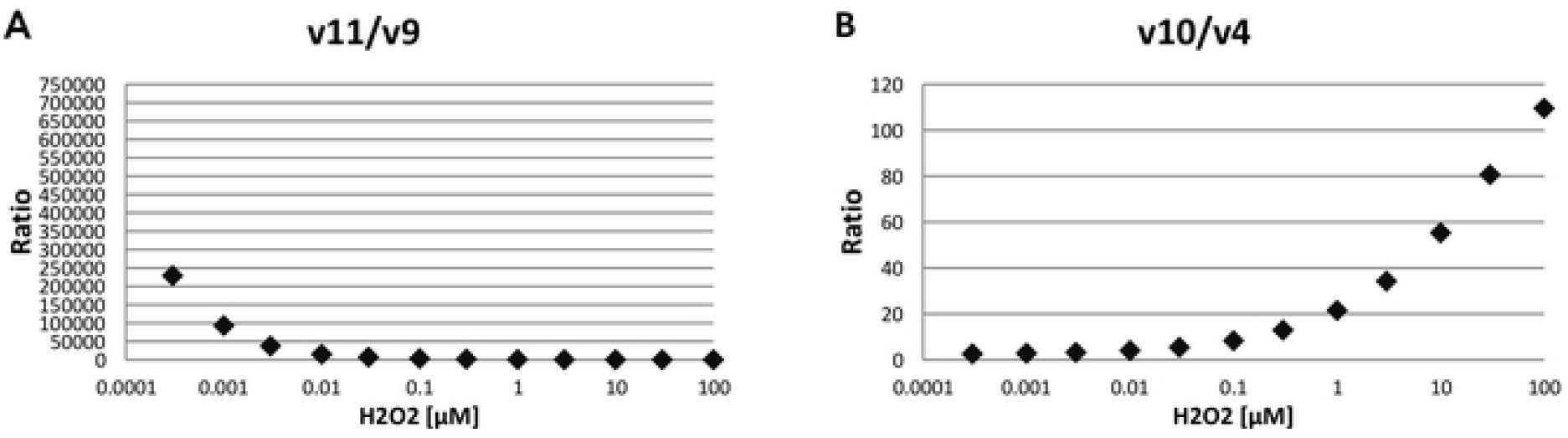
Steady state velocity ratios within the FTR/FNR network. Steady state velocities of the FTR/FNR-network (Fig. 5 A) were obtained after simulating the electron fluxes in the presence of various H_2_O_2_ concentrations. The physiological concentrations of network components were calculated for 1 mg chlorophyll. The H_2_O_2_ values were clamped in the simulation as given on the x-axis. (A) The ratio of the electron flux velocities from NADPH to metabolism (v11) relative to those from NADPH to thiol network (v9) were derived after 15 min. (B) Ratio of electron transfer rates from either TRX-f1 (v4) or NTRC (v10) to 2-CysPRX as a function of clamped H_2_O_2_ concentrations.

The third question dealt with the relative contribution of NTRC (v10) and TRX-f1 (v4) to reducing 2-CysPRX (Fig. 7). At low H_2_O_2_ concentrations v10 exceeded v4 by 2 to 3-fold; at 30 nM H_2_O_2_ the ratio of v10/v4 was 5.5. Apparently, the flux contribution of NTRC increased with increasing H_2_O_2_.

The final simulation explored the thermodynamic equilibrium between the NADPH system and the 2-CysPRX mediated by NTRC. The ratio of NADPH/NADP^+^ was varied between full reduction and full oxidation and the 2-CysPRX_red_/2-CysPRX_ox_ computed assuming full equilibrium catalyzed by NTRC (Fig. 8A). At a ratio of NADPH/NADP^+^=1only a small fraction of 2-CysPRX was in the oxidized form (Figs. 8A and B). Only at rather oxidized NADP system of 97.6% adjusted the 2-CysPRX system at the ratio of 35% reduced and 65% oxidized as reported in photosynthesizing leaves (Vaseghi et al. 2018). The computing result was confirmed experimentally with recombinant proteins of NTRC and 2-CysPRX equilibrated with varying NADPH/NADP^+^-ratios, labeled with 5 mM N-ethylmaleimide polyethylene glycol (mPEGmal) at pH 8 and separated on reducing sodium dodecylsulfate polyacrylamide gel electrophoresis (SDS-PAGE). The peroxidatic and resolving thiol of the reduced form bound two molecules of mPEGmal causing a shift of 10 kDa, while the disulfide bonded oxidized form could not be labeled and separated as a band at 24 kDa. In the presence of oxidized NADP^+^, only the oxidized form of 2-CysPRX was observed. The oxidized form decreased with increasing NADPH/NADP^+^-ratio. Importantly at a physiological NADPH/NADP^+^-ratio of 1, a significant amount of 2-CysPRX_ox_ was visible, albeit much less than 65% as reported [12].

**FIGURE 8:**
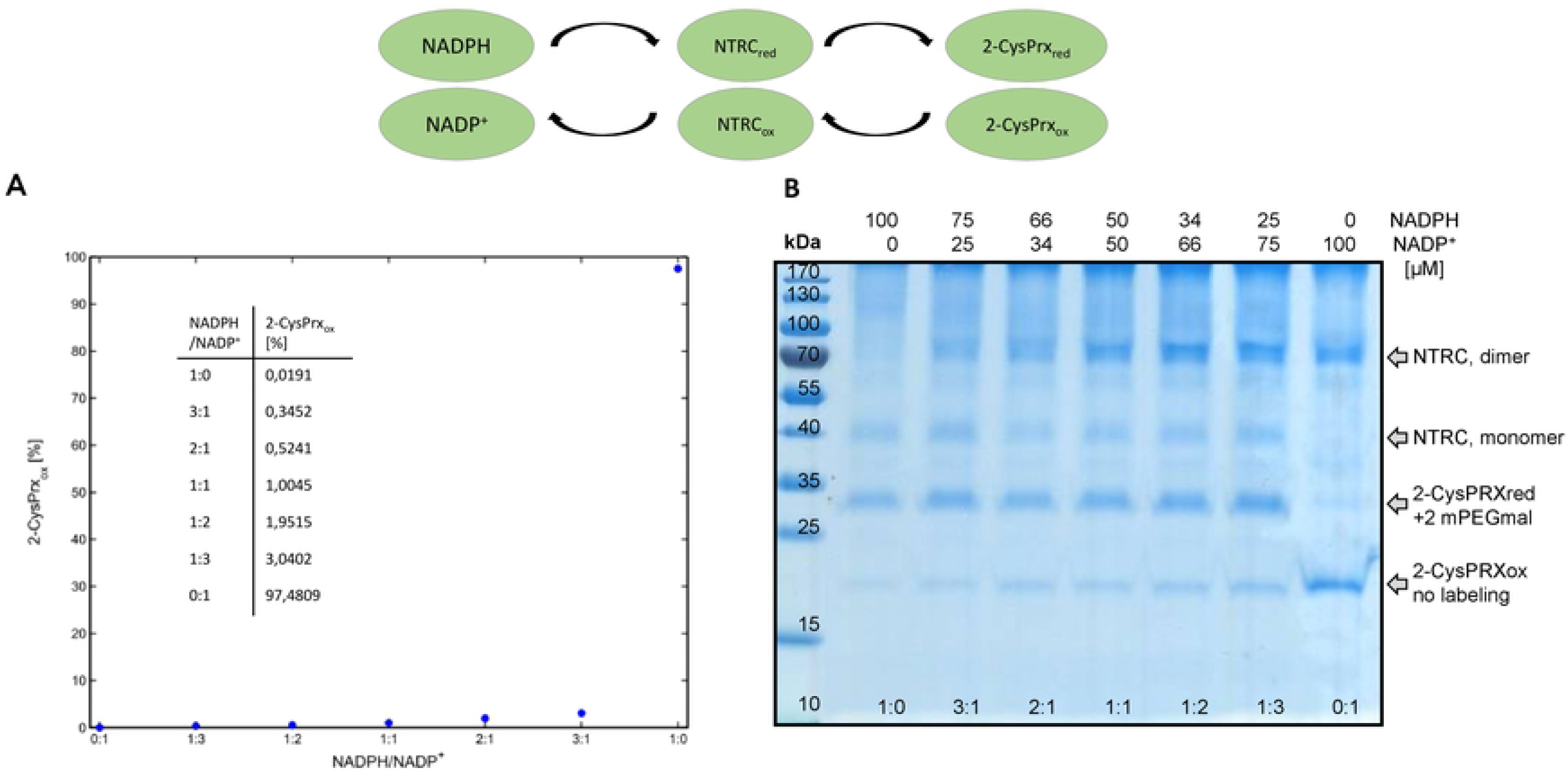
Redox equilibrium between NADPH and 2-CysPRX catalyzed by NTRC as computed *in silico* and measured experimentally in the reconstituted system. **(A) (B)** The equilibrium between varying NADPH/NADP^+^-ratios and 2-CysPRX was computed using a mathematical model consisting of differential equations. (B) Enzymatic assays containing 10 μM NTRC, 5 μM 2-CysPRX and 100 μM total (NADPH / NADP^+^) in TRIS-buffer, pH 8, were incubated for 5 min. After labeling the free thiols with mPEG-maleimide which causes an increase in molecular mass by 5 kDa per introduced label, thus 10 kDa for two thiols, samples were separated by SDS-PAGE and visualized by Coomassie-silver staining. The positions in the gel of the oxidized (no label) and reduced forms (two labels) of 2-CysPRX are indicated.

## DISCUSSION

Redox and reactive oxygen species-dependent signaling is a fundamental property of cells. For its understanding it is of fundamental importance to define the network connections, quantify electron fluxes and determine the driving forces [14]. Due to the network character, redox signaling can hardly be fully addressed experimentally. Thus work with mutants devoid of single and multiple network elements have provided important clues on their potential roles and functions, but also bear the problem of cumulating and equivocal effects [15,16]. For this reason, this study realized a computational approach for simulating two separate sub-networks and a combined network of the chloroplast as a meaningful approach complementary to the empiric avenue. In the following we will discuss the main conclusions drawn from our simulations and also address the potential shortcomings.

The FDX-FTR-branch was used to simulate the distribution of electrons between activation of an exemplary target protein, the chloroplast FBPase, and reduction of 2-CysPRX. The FBPase is only one of several targets of TRX-f1 [15]. *Arabidopsis thaliana* lacking TRX-f1 lacks an obvious phenotype. However the double mutant *ntrc/trx-f1* is compromised in multiple parameters such as growth, photosynthetic carbon assimilation and activation of FBPase. In parallel the increased NADPH/NADP-ratio in the double mutant indicates an inhibition of CBC activity [15]. But even the double mutant *trx-f1/trx-f2* showed a significant reduction of the FBPase and RubisCO activase protein, indicating alternative pathways for the reduction of photosynthesis-related target enzymes. But it is noteworthy that this simplified network allowed for simulating the data from the corresponding enzyme test surprisingly well [12]. The kinetic data of the network consisting of TRX-f1, FBPase and 2-CysPRX either reconstituted from recombinant proteins in a test tube or computed in silico matched with a regression coefficient of R^2^=0.998. This ‘perfect’ match confirms the reliability of these particular reaction constants.

The model cannot reflect the complexity of the chloroplast TRX system which consists of 20 TRX and TRX-like proteins (Trx-m (4) +Trx-x (1) + Trx-y (2) +Trx-f (2) + Trx-z (1) + Trx-Like2 (2) + Trx-Lilium (5) + CDSP32 (1) + HCF164 (1) + NTRC (1)) and the NTRC [3,17]. The implementation of additional TRXs in the model would require quantitative data on their stromal concentration and affinity toward targets of interest. However this information is unavailable for most chloroplast TRXs. It is an interesting perspective that such interactions may be predicted based in electrostatic and geometric properties of the complementary interfaces of redox transmitter and redox target in the future [18].

The FNR branch provides electrons from PET to NADPH which is mainly consumed in the CBC. NADPH also reduces NTRC. The reconstitution of NADPH/NTRC/2-CysPRX system showed the reversibility and equilibrium in this pathway. A highly oxidized NADP system oxidizes 2-CysPRX via NTRC. The data of Fig. 8 show that even in the presence of 75% oxidized NADP-system, only a small fraction of 2-Cys PRX turns oxidized. This result was in line with the theoretical computation of the redox equilibrium. Reverse flow from 2-CysPRX for NADP^+^ reduction will only occur if the NADP-system is oxidized to an overwhelming fraction which rarely occurs. Such a far-going oxidation of the NADPH/NADP^+^-ratio was reported for spinach leaves when lowering the steady state light intensity from 250 μmol photonsm^−2^s^−1^ to 25 μmol photonsm^−2^s^−1^ [19]. Thus the backflow may be a feedback mechanism upon sudden lowering or extinguishing the photosynthetic active radiation. After such a light step down, the CBC still consumes NADPH and strongly oxidizes the NADP-system, which oxidizes the 2-CysPRX by backflow. This mechanism will accelerate the TRX oxidation by 2-CysPRX acting as TRX oxidase [12] and thereby downregulates the CBC activity to readjust the NADPH/NADP^+^-ratio to reach an energetic equilibrium.

Simulating the effect of H_2_O_2_ using the model combined from the FTR and FNR networks allowed for estimating velocities of empirically inaccessible reactions and amounts of resting H_2_O_2_ concentrations. Biochemical H_2_O_2_ determination in extracts or histochemical staining only provide rough estimates and possibly indications for alterations, but these quantifications give unrealistically high ROS amounts. Recent developments with H_2_O_2_-sensitive *in vivo* probes such as Hyper enable kinetic monitoring of H_2_O_2_ amounts in compartments of living cells. HyPer2 is a derivative of YFP fused to the H_2_O_2_ binding domain of the bacterial H_2_O_2_-sensitive transcription factor OxyR [20]. Using this sensor, Exposito-Rodriguez et al. [21] proved that chloroplast-sourced H_2_O_2_ likely are transported to the nucleus. The study exclusively was based on excitation ratios but H_2_O_2_ concentrations could not be estimated.

The steady state concentration of stromal H_2_O_2_ was approximated to about 30 nM in this study. The rational was to compare the electron distribution and computed redox states of network components in the presence of different H_2_O_2_ concentrations with reported data on the redox state of 2-CysPRX *ex vivo* [9,12]. The high reaction rate of 2-CysPRX with peroxide substrates [22] allows for rapid oxidation of the peroxidatic thiol and conversion to the disulfide form [23]. The limiting factor in the catalytic cycle is the regeneration [13,24]. The limited regeneration speed decreases the turnover number to values far below 1 s^−1^. Consequently, any increase in H_2_O_2_ will shift the 2-CysPRX redox state to more oxidation. The value of 30 nM could be an underestimation if other TRX isoforms or other electron donors significantly contribute to the reduction of disulfide-bonded 2-CysPRX. The most interesting candidate is TRX-x which proved to be the most efficient regenerator of 2-CysPRX among the tested TRXs [7], but had little effect on the redox state of 2-CysPRX measured *ex vivo* (Pulido et al. 2010). This may not be surprising since the fraction of TRX-x only amounts to 8% of that of TRX-f1 and 5% of that of NTRC in the stroma according to the AT_CHLORO mass-spectrometric protein database [25,26].

Electrons from light-driven PET are distributed among different metabolic consumers such as carbon, nitrogen and sulfur assimilation which are serviced at a ratio of about 40:8:1. In addition part of the electrons are used for regulatory purposes, namely for producing both the reductants NADPH, glutathione and TRX as redox input elements into the thiol redox regulatory network [27] and the oxidant H_2_O_2_ [28]. The relative expenditure of reductive energy for redox regulation of the CBC cycle has been an open but unsolved issue for long, essentially since the discovery of TRXs. The mathematical simulation focusing on FBPase assumed that both the reductive and the oxidative driving forces are generated from PET. In this case and at a resting H_2_O_2_ concentration of 30 nM, metabolic electron drainage exceeds the NTRC-dependent regeneration of 2-CysPRX by a factor of 7234-fold (Suppl. Table 4). Including the 5.46-fold lower electron flux at 30 nM H_2_O_2_ from TRX-f1 for 2-CysPRX regeneration (Suppl. Table 4) the metabolic flux exceeds the reduction rate of 2-CysPRX 6114-fold. An equivalent amount of electrons must be used to produce H_2_O_2_, increasing the reductive expenditure for FBPase redox regulation to 1/3057^th^ of metabolic flux. Considering the other redox regulated targets such as RubiCO activase, seduheptulose-1,7-bisphosphatase, glyceraldehyde-3-phosphate dehydrogenase, ribulo-5-phosphate kinase and malate dehydrogenase and assuming that regulation of these targets consumes, e.g., 30-fold more electrons than regulation of FBPase, then about 1% of the PET rate would be drained for redox regulation.

Another unknown parameter in the system is the nature of oxidation in addition of PET-derived H_2_O_2_. Two sources for oxidation should be taken into account. H_2_O_2_ is produced outside of the chloroplast, in particular in the peroxisomes, in mitochondria and at the plasmamembrane by NADPH oxidases [29]. Antioxidant systems decompose these ROS and thus it is unlikely that external H_2_O_2_ penetrates the chloroplast and contributes to oxidation of redox target proteins. Another possible oxidant is elemental oxygen as suggested early after the discovery of thioredoxins. It would be important to obtain the kinetic data of O_2_-mediated oxidation of TRX and other protein thiols in future work in order to incorporate such data in the mathematical model. Alternative oxidation reactions will increase the expenditure of electron for regulation.

A unique model simulating the reactive oxygen species network of the chloroplast was constructed by Polle in 2001 [30]. It focused on the water-water cycle but did not include PRX and redox regulation. The simulation showed that neither O_2_^−^ nor H_2_O_2_ accumulate in the chloroplast as long as the supply with reductants is maintained high. The H_2_O_2_ concentration was estimated in the submicromolar range. The focus here was placed in redox regulation of a CBC target protein. Additionally, the redox state of the 2-CysPRX provided a benchmark for estimating the resting H_2_O_2_ concentration. Thus the H_2_O_2_ production and detoxification rates establishing this low H_2_O_2_ concentration are not of primary importance for our model.

## MATERIALS AND METHODS

### Equilibrium between NADP-system and 2-CysPRX catalyzed by NTRC

Hisx6-tagged recombinant NTRC and 2-CysPRX were produced in *E. coli* and purified by Ni-nitrilotriacetic acid-based affinity chromatography as described [12]. 10 μM recombinant NTRC was incubated with 5 μM of 2-CysPRXA and 100 μM NADPH/NADP^+^ in 50 mM Tris-HCl pH 8 in a final volume of 50 μl for 5 min. Then 50 μl of TCA 20% (w/v) was added to the mixture and maintained on ice for 40 min. The assay mix was spun for 15 min at 13,000 rpm. The pellet was washed with TCA (2%, 100 μl). After 15 min centrifugation at 13 000 rpm, the pellet was resuspended with 15 μl of 50 mM Tris-HCl pH 7.9 containing mPEGmal with 1% SDS. After 90 min at room temperature, SDS-PAGE loading buffer with ß-mercaptoethanol was added. 20 μl of the mixture were separated by SDS-PAGE (12% w/v) and protein bands visualized with Coomassie-silver staining.

### Concentration of network components

Concentration of chloroplast proteins were taken from literature and calculated for 1 mg Chl refered to 66 μl stroma [31] and 10 mg stromal protein. The calculated concentration values were summed for isoforms. In all models each H_2_O_2_ and FDX concentration were set constant. The start values of variables were partitioned into 80 % of reduced and 20 % of oxidized form except for NTRC, 2-CysPRX and NADPH/NADP^+^ couple (Suppl. Table 5).

### Model formulation

Three chloroplast network models were developed to analyze electron transfer rates as well as oxidized and reduced states of network components with various H_2_O_2_ concentrations. The first model describes the FTR-based electron transfer to 2-CysPRX (Fig. 1A), the second model reveals the FNR-based electron transfer (Fig. 3A) and the third model combines both models (Fig. 5A).

#### A) FTR network model

In order to analyse the electron distribution from TRX-f1 to FBPase or 2-CysPRX in dependence on H_2_O_2_ concentration, the first simplified model of the FTR network consisted of FDX, FTR, TRX-f1, FBPase, 2-CysPRX and H_2_O_2_ (Fig.1 A). FDX and H_2_O_2_ were constant parameters. FDX was constantly reduced by 50 % and the H_2_O_2_ concentration varied from 0.3 nM to 10 μM. The four variables were FTR, TRX-f1, FBPase and 2-CysPRX. Each variable could adopt the oxidized and reduced state. Electrons were transferred from FDX (v1) via FTR to TRX-f1 (v2). TRX-f1 distributed the electrons to FBPase (v3) and 2-CysPRX (v4). H_2_O_2_ oxidized 2-CysPRX (v5). The rate equations were implemented using mass action law (Appendix A1). In general the reactions were formulated as reversible second order rates.

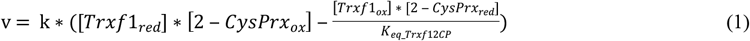

The transition from one electron transfer to two electron transfer takes place at FTR. Therefore, two FDX are required to reduce FTR.

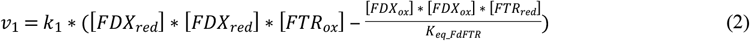

#### B) FNR network model

The second model aimed to describing the reduction power in the FNR branch toward 2-CysPRX and represents the electron transfer via FNR and NTRC (Fig. 3A). This FNR network model consisted of FDX, FNR, NADPH, NTRC, 2-CysPRX and H_2_O_2_. Each component exhibited two states in the model; oxidized and reduced form. Only FNR (the transition molecule from one to two electron transport) was represented in three forms; reduced, half reduced and oxidized. Electrons were transferred from FDX (constant reduced 50%) to FNRox (v1) that results in half reduced FNR form.

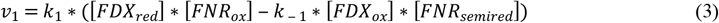

A further reduction by FDX of FNRsemired (v2) resulted in the fully reduced form of FNR (FNRred).

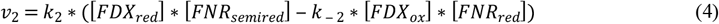

FNRred transfered electrons to NADP^+^ (v3) to produce NADPH. In order to mimic metabolic NADPH consumption an estimated rate of NADPH decrease was included (v7). In this network NADPH transfered electrons to NTRCox (v4) that led to reduced NTRC (NTRCred). The reduction of 2-CysPRXox took place by NTRCred (v5). Reduced 2-CysPRX reduced H_2_O_2_ to 2 H_2_O (v6). H_2_O_2_ was included as a constant and varied from 0.3 nM to 100 μM. (Appendix B)

#### C) The combined FTR-FNR network model

To analyse the interaction between the FTR and FNR branch in adjusting the 2-CysPRX and FBPase redox states in dependence on different H_2_O_2_ concentrations a third model was constructed consisting of the FTR and FNR networks (Fig. 5 A). All components except FDX and H_2_O_2_ were variables and represented in reduced and oxidized form. Only FNR adopted three different redox states; reduced, oxidized and half reduced.

The equilibrium constants *K_eq_* in all reactions were calculated using the standard cell potentials *E^o^* of each cell reaction at pH 7 linked to the standard reaction Gibbs energy *Δ_R_G*° [32]:

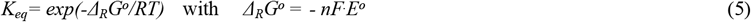

where *R* is the gas constant, *T* the thermodynamic temperature, *F* the Faraday constant and *n* the stoichiometric coefficient of the electrons in the half-reactions in which the cell reaction can be divided. The models were formalized as systems of differential equations (Appendix C). Steady-state solutions were computed numerically in MATLAB.

### Fitting of unknown parameter

Unknown parameters were fitted to data [33]. A model was developed containing all assay components. The fitting procedure employed genetic algorithms and root mean squares for comparison of fitting quality in MATLAB. The network consisted of NADPH (0.5 mM), FNR (0.2 μM), FDX (1 μM), FTR (1μM), TRX-f1 (2 μM) (Suppl. Fig. 4).

## Appendix A Model equation

To represent the thiol-disulfide redox network of the cell three different models were constructed. The FTR model (Fig. 1A), the FNR model (Fig. 2A) and after their merging the FTR-FNR model (Fig. 3A).

### A1 FTR network model

In the model represented in Fig. 1A. FDX and H_2_O_2_ are external quantities and their concentration is constant. The variables FTR, TRX-f1, FBPase and 2-CysPRX exhibit a reduced and oxidized form (reaction equation Suppl Table 6). The rate expressions are read

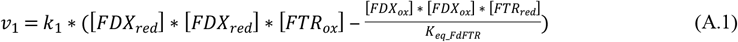

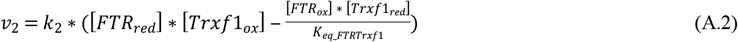

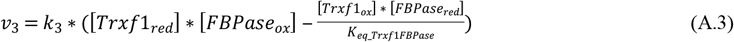

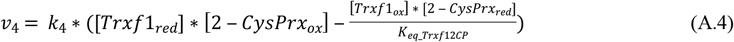

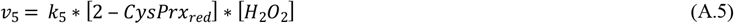

FDX is implemented with a constant redox state of 50 % reduced and oxidized. Constant H_2_O_2_ concentration vary form 0 to 10 μM in different simulation.

The differential equations of the variables used for simulating FTR network model are:

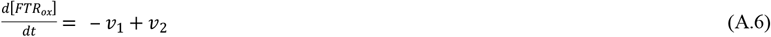

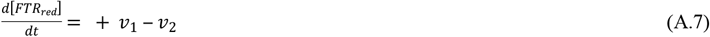

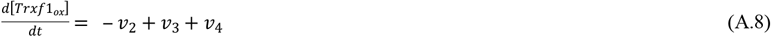

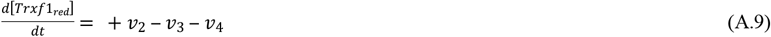

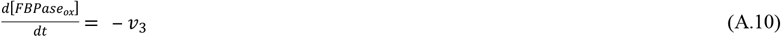

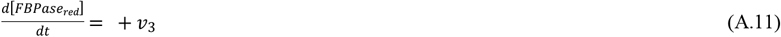

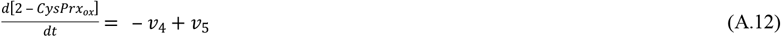

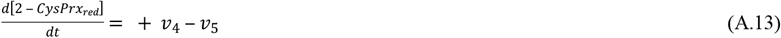

### A2 FNR network model

The FNR network model (Fig 3A) consists of FDX, FNR, NADPH, NTRC, 2-CysPRX and H_2_O_2_. Each component has an oxidized and reduced form. Only FNR exhibits three forms; reduced, half reduced and oxidized. In order to mimic metabolic NADPH consumption an estimated rate of NADPH decrease was included (v7). Each reaction (except v7) is reversible. The equilibrium constants are calculated from redox potential of involved components (material and methods). The rate expressions of FNR network model are

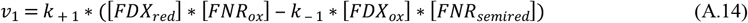

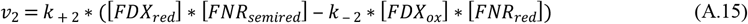

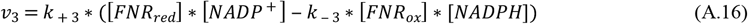

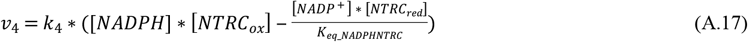

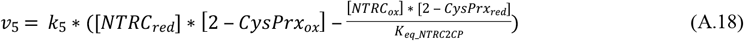

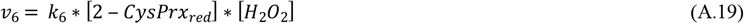

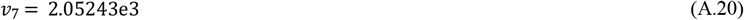

FDX and H_2_O_2_ concentrations are considered to be constants. FDX is constantly reduced at 50 % and the H_2_O_2_ concentration varies from 0 to 100 μM in different simulations.

The differential equation of the variables used for simulating FNR network model are:

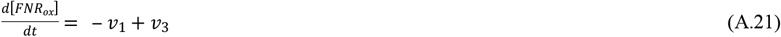

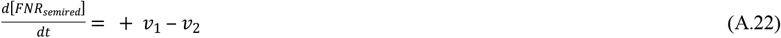

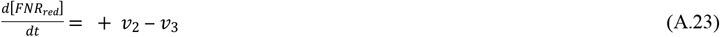

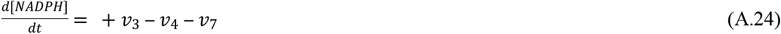

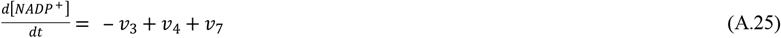

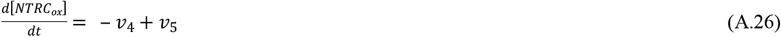

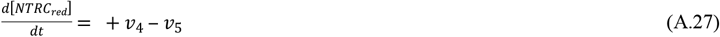

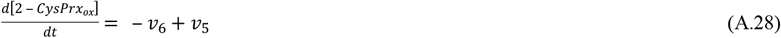

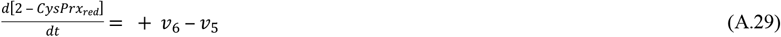

### A3 FTR-FNR network model

The FTR-FNR network model (Fig. 5A) combines both submodels. FDX and H_2_O_2_ are considered to be constant quantities. The redox state of FDX is set constant to 50 % reduced form and the concentration of H_2_O_2_ varies from 0.3 nM to 100 μM. The variables are FTR, TRX-f1, FBPase, 2-CysPrx, FNR, NADPH/NADP^+^ couple and NTRC. Each variable (except FNR) is represented in an oxidized and reduced state. FNR is implemented in three forms, oxidized, reduced and half reduced. Each reaction (except the metabolic consumption of NADPH; v11) is reversible. The equilibrium constants are calculated from redox potentials of involved components (see Material and Methods). The rate expressions of FTR-FNR network model are

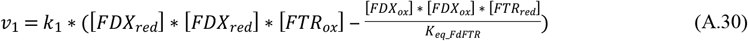

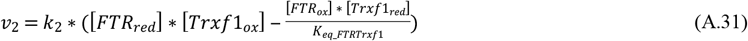

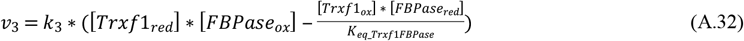

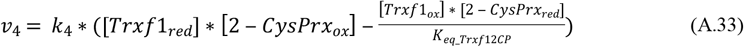

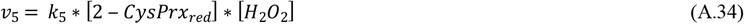

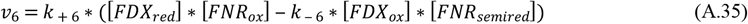

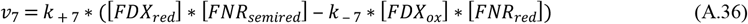

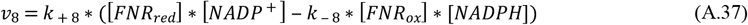

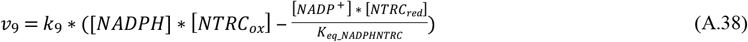

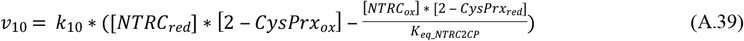

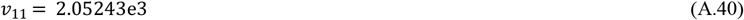

The complete differential equation system introduced into MATLAB is the following:

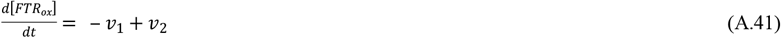

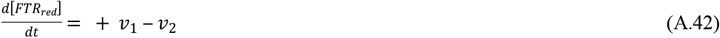

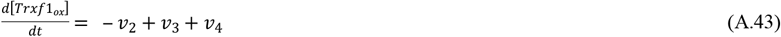

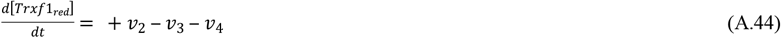

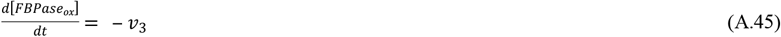

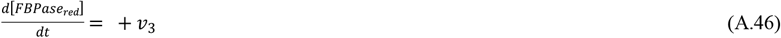

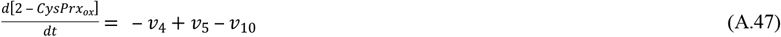

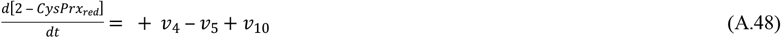

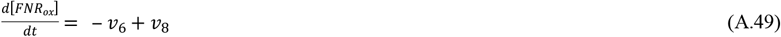

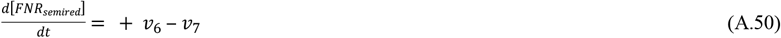

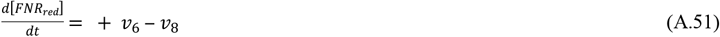

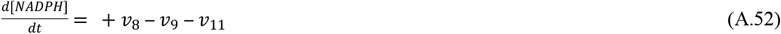

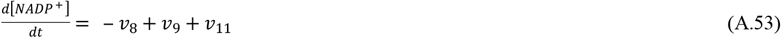

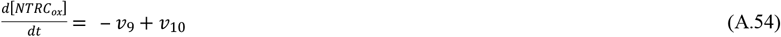

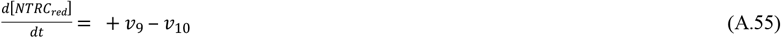

## Appendix B Choice of parameter

The parameters of FTR-FNR network model are given in Table B.1. Most of the parameters are available from literature. The units of concentrations are μM/s. The rate constants are second or third order. Unknown rate constants are fitted (see Material and Methods). The physiological concentrations of network components are calculated for 1 μg Chl and 66 μL stromal volume (Winter *et al.*, 1994). If needed, the concentrations of isoforms are summed up.

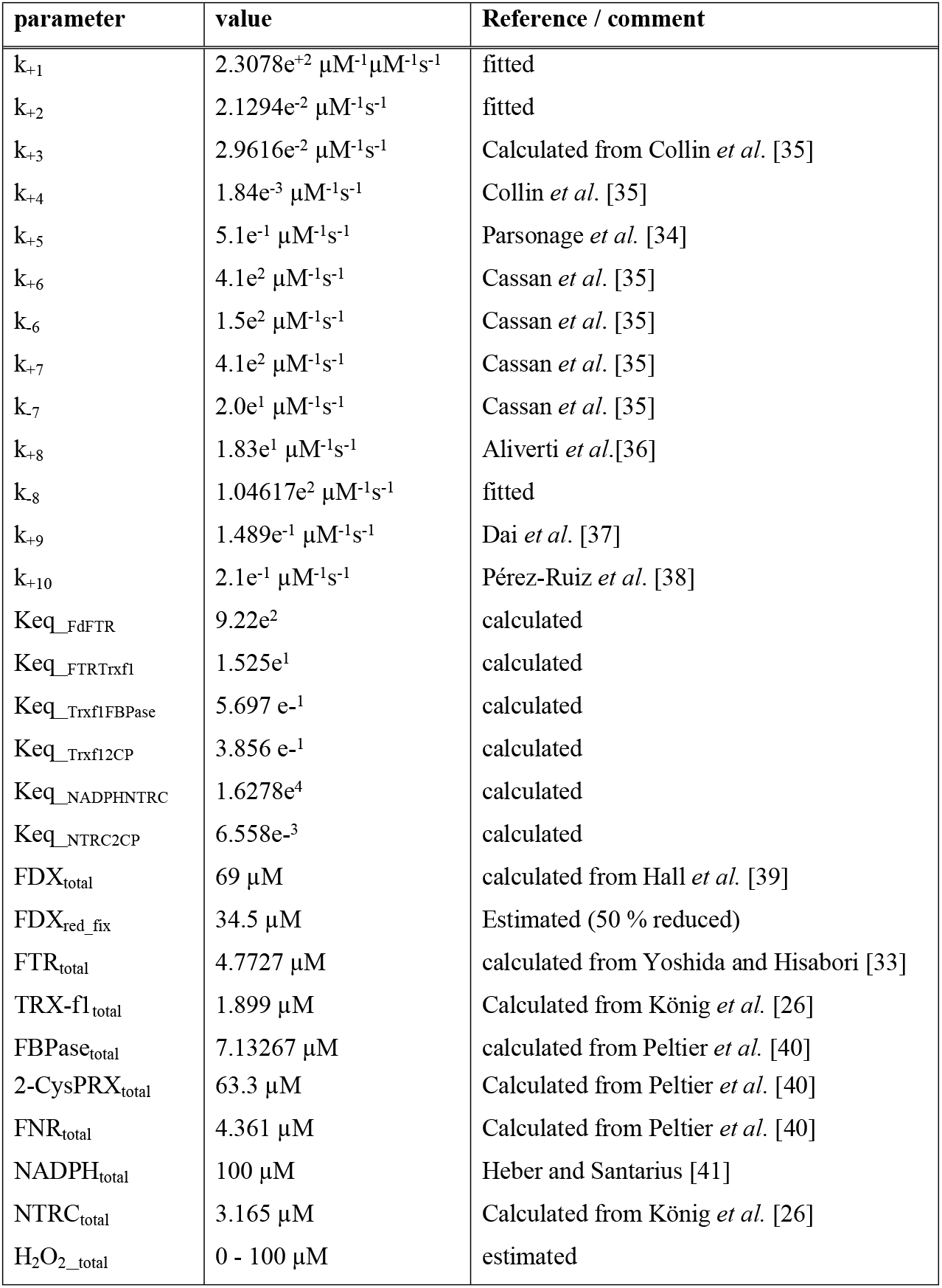

## AUTHORS CONTRIBUTION

MG designed the study, implemented the model, wrote the paper; SK supported the implementation of the mathematical model; KC performed the equilibration experiment between varying NADPH/NADP^+^-ratios, NTRC and 2-CysPRX; KJD designed the study, discussed the results and wrote the paper.

## ACKNOWLEDGEMENTS

The financial support of the own work by Bielefeld University and the DFG (Di 346-14, -17) is gratefully acknowledged.

## SUPPLEMENTARY MATERIALS

**Suppl. Figure 1**: Simulated steady state of FNR-network components in dependence on H_2_O_2_ concentrations in the absence of electron drainage by metabolism.

**Suppl. Figure 2**: Simulation of time-dependent redox potential changes of FNR-network components in the absence of metabolic electron drainage.

**Suppl. Figure 3**: Model comparison by simulation of model components over time.

**Suppl. Figure 4**: Fitting of unknown parameter.

**Suppl. Table 1**. Simulated steady state concentrations of FTR-network components.

**Suppl. Table 2**. Simulated steady redox state of the FNR-network components.

**Suppl. Table 3**. Simulated steady state concentrations of the FTR/FNR-network components.

**Suppl. Table 4**. Calculated ratios of steady state velocities observed in the combined FTR/FNR-network.

**Suppl. Table 5**. Distribution of network components in reduced and oxidized form at t=0 in FTR-FNR model.

**Suppl. Table 6**. Reaction equations describing the model of FTR network model.

**Suppl. Table 7**. Reaction equations describing the model of FNR network model.

**Suppl. Table 8**. Reaction equations describing the model of the combined FTR-FNR network model.

**Suppl. Table 9**. Redox potentials of network components.

**Suppl. Table 10**. Parameters of FTR network model.

**Suppl. Table 11**. Parameter of FNR network model.

